# Rapid inference of direct interactions in large-scale ecological networks from heterogeneous microbial sequencing data

**DOI:** 10.1101/390195

**Authors:** Janko Tackmann, João Frederico Matias Rodrigues, Christian von Mering

## Abstract

The recent explosion of metagenomic sequencing data opens the door towards the modeling of microbial ecosystems in unprecedented detail. In particular, co-occurrence based prediction of ecological interactions could strongly benefit from this development. However, current methods fall short on several fronts: univariate tools do not distinguish between direct and indirect interactions, resulting in excessive false positives, while approaches with better resolution are so far computationally highly limited. Furthermore, confounding variables typical for cross-study data sets are rarely addressed. We present FlashWeave, a new approach based on a flexible Probabilistic Graphical Models framework to infer highly resolved direct microbial interactions from massive heterogeneous microbial abundance data sets with seamless integration of metadata. On a variety of benchmarks, FlashWeave outperforms state-of-the-art methods by several orders of magnitude in terms of speed while generally providing increased accuracy. We apply FlashWeave to a cross-study data set of 69 818 publicly available human gut samples, resulting in one of the largest and most diverse models of microbial interactions in the human gut to date.

## Introduction

Microorganisms shape virtually every aspect of Earth’s biosphere. Besides their critical role in global geochemical cycles [1] and widespread symbiosis with all major branches of life [2–4], the tight coupling between the microbiome and human health is rapidly gaining appreciation [5, 6]. While the structure of microbial ecosystems is influenced by environmental factors and hosts [7–9], another important driving force are ecological interactions between microbes [10, 11], such as competition, symbiosis, commensalism, and antagonism.

The inability to (co-)culture the majority of microorganisms in the lab [12, 13] makes computational tools that predict ecological dependencies between microbes pivotal. Common to these approaches is the exploitation of cross-sectional (co-occurrence and co-abundance [14, 15]) and temporal [16, 17] statistical patterns, or metabolic complementarity [18, 19], to infer ecological associations and construct interaction networks. While currently widespread methods are mostly restricted to pairwise interactions based on univariate statistical associations [17, 20], more recent approaches based on Probabilistic Graphical Models (PGM) consider the conditional dependency structure between microbes and thereby allow for the distinction between direct and indirect interactions [15, 21]. While approaches of this class result in reduced false positive edges and thereby infer more sparse and interpretable networks, they usually require larger data sets with sufficient statistical power. Hundreds of thousands of microbial sequencing samples from varying environments and studies all around the globe are now available [22, 23], yet this rich vault of data can currently not be utilized as state-of-the-art PGM methods for ecological inference are unable to computationally scale to these dimensions. Furthermore, particular properties of these large meta-study data sets, such as variation in habitats, measurement conditions and sequencing technology can lead to high amounts of confounded associations, which current methods typically don’t address.

Here, we present FlashWeave, a novel approach for inference of high-resolution interaction networks from large and heterogeneous collections of microbial sequencing samples based on co-occurrence. Flashweave is highly optimized for computational speed and addresses a number of known artifacts common in large cross-study sequencing data sets, such as compositionality effects, bystander effects, shared-niche biases and technological biases. It furthermore allows the seamless integration of environmental factors, such as temperature and pH, which enables estimation of their influence on the studied ecosystem and removal of interactions they drive, paving the path towards microbial systems biology at a global scale.

We compare FlashWeave to a variety of state-of-the-art methods on a wide collection of synthetic and biological benchmark data sets and show that FlashWeave markedly outperforms other methods in terms of speed, while typically achieving comparable or increased accuracy, in particular on heterogeneous data sets with large fractions of structural zeros. We furthermore illustrate the usefulness of integrating environmental and technical factors through inclusion of habitat and primer variables into the inference of an interaction network based on the Human Microbiome Project. Finally, we apply FlashWeave to a global collection of 69 818 publicly available metagenomic sequencing samples from the human gastrointestinal tract, covering 488 projects, and infer the to date largest interaction network of the human gut using minimal computational resources and time. The network is consistent with previously described biological patterns and additionally features many connections within the to date underexplored rare biosphere. It further unveils important candidates for novel mutualist and negative hubs that warrant further investigation and yields an unusually strong signal for phylogenetic assortativity, pointing towards pronounced kin selection.

## Results

### A fast and compositionally robust method for ecological network inference, with handling of heterogeneous data

FlashWeave is based on the Local-to-Global Learning (LGL) approach proposed by Aliferis et al. [24], a constraint-based causal inference framework for prediction of direct relationships between variables in large systems. Algorithms of this family infer the Markov blanket of each target variable *T*, which constitutes the directly associated neighborhood *MB(T*) rendering all remaining variables *S* probabilistically independent of *T*. It thus removes spurious, i.e. purely correlational, associations commonly reported by wide-spread univariate methods. Related algorithms have been successfully applied in a wide range of fields, including cancer diagnosis [25], drug-drug interactions [26] and gene regulatory network inference [27].

FlashWeave is a highly optimized implementation of the semi-interleaved HITON-PC [24] instantiation of LGL (Fig. 1 A), critically extended with several high-performance heuristics (see Text S1), as well as methods addressing heterogeneous data and state-of-the-art compositionality correction (see Text S1). The latter is essential since abundances from sequencing data constitute compositions and are thereby constrained to the simplex, which has long been known to induce artificial correlations [28, 29].

**Fig. 1.**
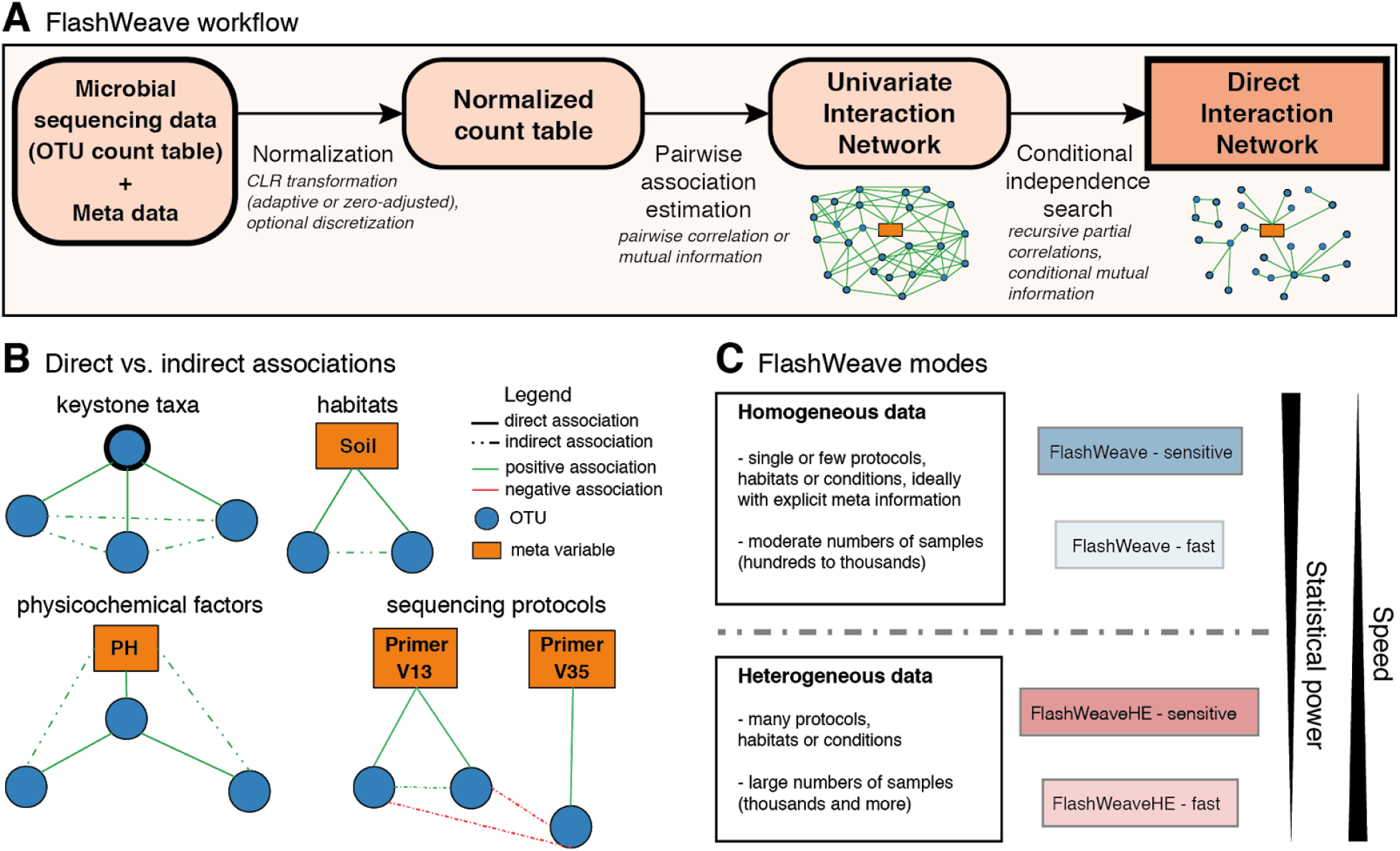
Overview of FlashWeave. **A** Main steps in the network inference pipeline. **B** Examples of how indirect associations may create false positive results in a variety of ecological and technological scenarios. **C** Use cases for the different FlashWeave modes.

In contrast to most other methods, FlashWeave can furthermore utilize meta variable (MV) information (Fig. 1 B), such as subject lifestyle factors, physicochemical measurements or sequencing protocol information to i) further reduce spurious associations and ii) report direct OTU-MV relationships.

### Prediction performance on a variety of synthetic data sets

Since experimentally verified biological interactions between microbes are scarcely available, we first employed previously published frameworks that generate synthetic data with ecological structure. We compared the quality of networks inferred by FlashWeave “sensitive” (-S) and “fast” (−F) (Fig. 1 C) to three competing univariate inference methods (SparCC [30], eLSA [17] and CoNet [20]) and three conditional methods (mLDM [21] and SpiecEasi [15] with neighborhood selection (MB) and inverse covariance selection (GL)).

The first group of benchmark data sets was generated with a method from [15] based on the Normal to Anything (NorTA) approach. It takes abundance data from sequencing experiments and a custom interaction graph as an input and generates data with OTU distributions fitted to a target distribution, respecting constraints in the experimental data and the partial correlations from the input graph.

Along a gradient of increasing data set sizes fitted to a filtered version of the American Gut Project [31] data set, FlashWeave most accurately reconstructed the input graphs in most cases as measured by F1 scores of predicted edges (Fig. 2 C). Across topologies, non-FlashWeave methods achieved average F1 scores of 9% (CoNet) to 89% (SpiecEasi-MB) relative to FlashWeave-S (mean across methods: 52%). FlashWeave-S performed particularly well on the scale-free topology, where the closest method, SpiecEasi-MB, achieved 27% smaller F1 scores on average. FlashWeave-F was generally less predictive than FlashWeave-S with mean F1 score reductions of 2% (cluster) and 24% (scale-free).

**Fig. 2.**
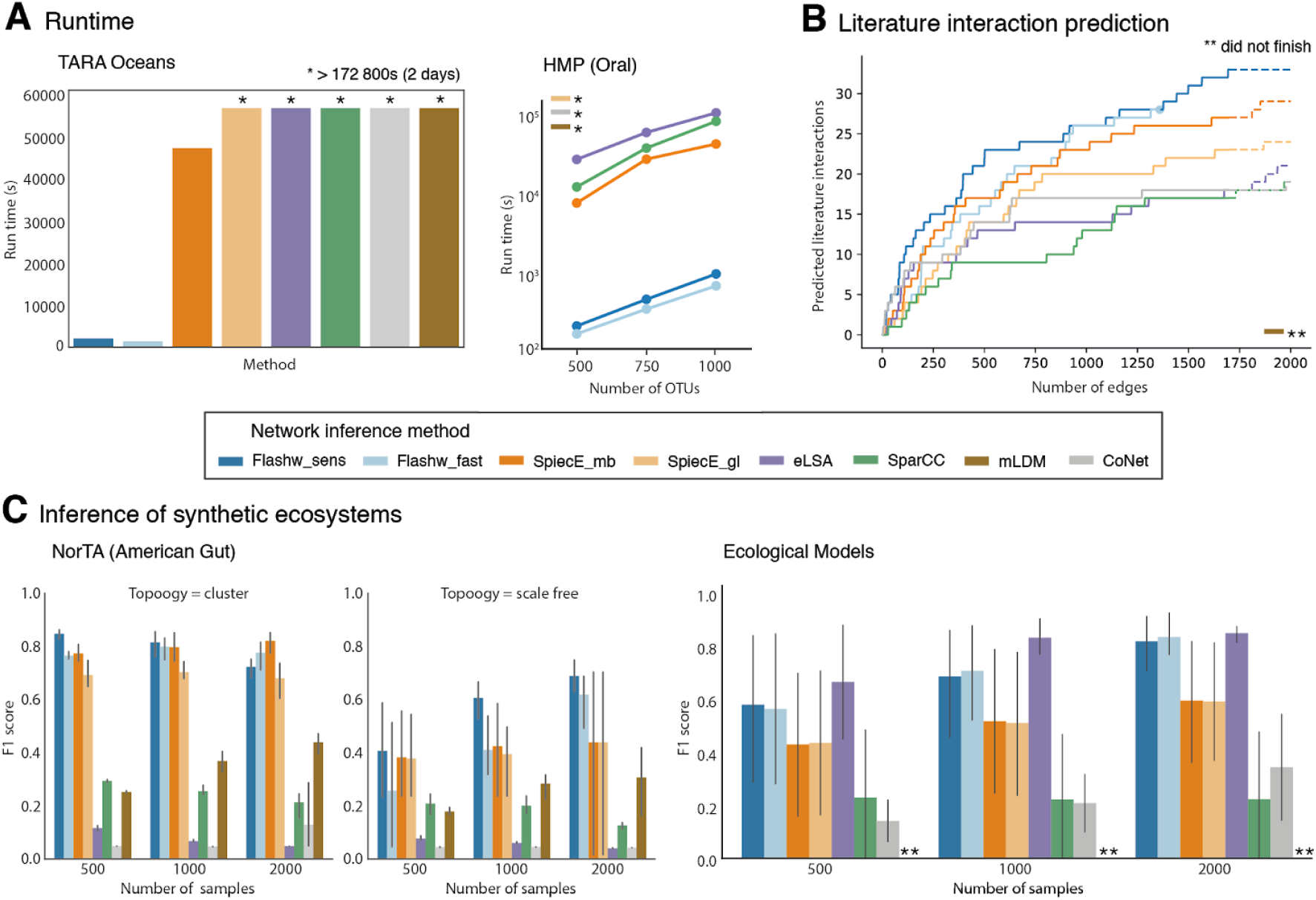
Comparison of FlashWeave to competing state-of-the-art network inference methods. Method abbreviations are Flashw_sens: FlashWeave-S, Flashw_fast: FlashWeave-F, SpiecE_mb: SpiecEasi-MB, SpiecE_gl: SpiecEasi-GL. **A** Run time comparison on the TARA Oceans and Human Microbiome Project (oral body site only) data sets. **B** Ranking of gold-standard planktonic interactions inferred from literature in the TARA Oceans data set among the top 2000 edges predicted by each tool. mLDM did not finish computation after two weeks. **C** Prediction performance on data sets with synthetically engineered edges, data generated based on[15] and[32], measured as F1 score (harmonic mean of precision and recall). Error bars depict 95% confidence intervals of the mean, based on 1000 bootstrap replicates.

In a second accuracy benchmark (“Ecological Models”) we used methods presented in [32] to generate abundance tables with a wide range of linear ecological relationships between OTUs, featuring varying degrees of sparsity and compositionality (Fig. 2 C). Across all data set sizes, eLSA achieved the highest F1 scores (mean 0.76), followed by FlashWeave-S (mean 0.68, 90% of eLSA). Notably, FlashWeave-S scores were almost identical to eLSA at the highest number of samples (99% median). FlashWeave-S and FlashWeave-F showed comparable results (< 2% difference), while all other methods achieved mean F1 scores of 2% (SparCC) to 73% (SpiecEasi-MB) compared to FlashWeave-F. In both the NorTA and the Ecological Models benchmarks, FlashWeave predictions generally improved with higher sample numbers (up to 141%), indicating efficient usage of additional data.

In order to test edge accuracy of FlashWeaveHE, which specializes on heterogeneous data (Fig. 1 C), we increased heterogeneity in the testing data by treating the three data sets per ecological table from the Ecological Models benchmark as separate, disjoint habitats and aggregating them into a single data set per table. FlashWeaveHE-S achieved the highest F1 scores on this benchmark (mean 0.78, Fig. 3 D), followed by FlashWeaveHE-F with 0.6 and FlashWeave-F with 0.43. The best non-FlashWeave method, SpiecEasi-GL, achieved a median of 0.25, 68% less than FlashWeaveHE-S. Notably, FlashWeaveHE modes displayed almost perfect precision (0.99) while non-FlashWeave methods ranged from 0.0007 (SparCC) to 0.2 (SpiecEasi-GL).

**Fig. 3.**
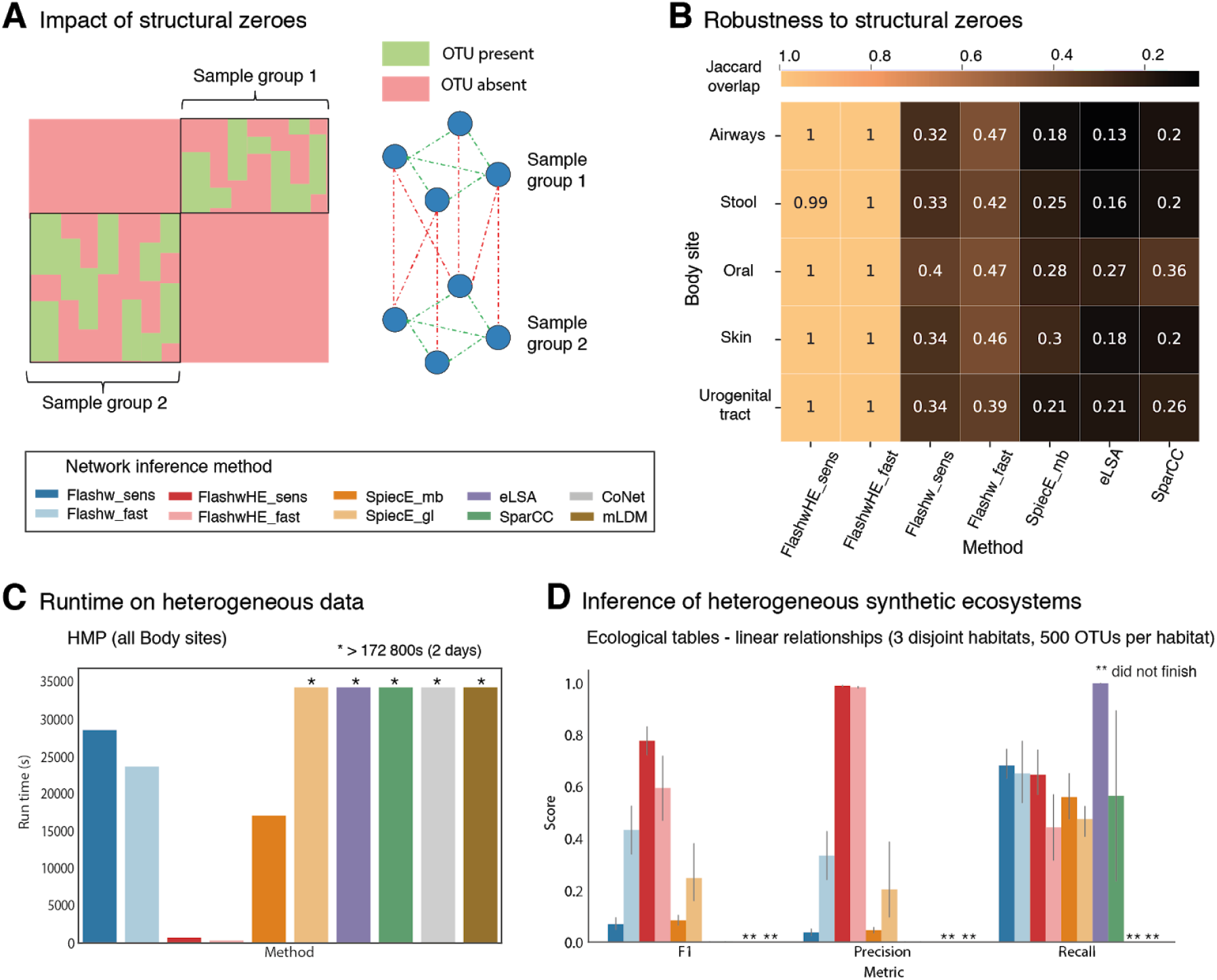
Network inference performance on heterogeneous data sets with increased structural sparsity. Method abbreviations (in addition to Fig. 2) are FlashwHE_sens: FlashWeaveHE-S, FlashwHE_fast: FlashWeaveHE-F. **A** Overview of how structural zeros can lead to false positive edges. **B** Similarity of HMP sub-networks computed on i) OTUs from a single body site (no structural zeros) or ii) body-site specific OTUs from all sites combined (structural zeros). SpiecEasi-GL, CoNet and mLDM did not finish computation after two weeks. **C** Run time comparison on the HMP data set (all body sites) as a representative heterogeneous data set. **D** Prediction performance on aggregated disjoint habitats generated by the Ecological Models approach[32] measured using different metrics. CoNet and mLDM did not finish computation after two weeks. Error bars depict 95% confidence intervals of the mean, based on 1000 bootstrap replicates.

### Reconstruction of literature interactions in TARA Oceans

In the original study of planktonic associations in the TARA Oceans project, the authors presented a list of gold-standard genus-level interactions described in the literature. This set provides an approximation of ground-truth against which network inference tools can be tested. Notably, it only comprises interactions between a small fraction of the total marine micro-eukaryotic diversity and is thereby likely incomplete. It thus allows only the comparison of a subset of true positives and precludes quantification of false positives. Consequently, methods that predict more edges are a priori at an advantage when raw numbers of true positives are compared, since such tools may have increased sensitivity but also higher false-positive rates.

In order to attempt a fair benchmark, we therefore compared methods in terms of how highly they rank literature interactions amongst their 2000 strongest reported associations, assuming that methods giving more weight to known interactions tend generally report more reliable relationships. In order to allow all methods to finish computation, we reduced the TARA Oceans data set to only OTUs that participate in at least one literature interaction.

FlashWeave-S found on average 24% more literature interactions among high-ranking edges than the closest follow-up method (SpiecEasi-MB), 38% more than FlashWeave-F and 80% more than other methods. While the TARA Oceans data set shows considerable heterogeneity, FlashWeaveHE was not applicable due to insufficient statistical power (only 22 - 335 predicted edges total).

### Speed comparison on the Human Microbiome Project and TARA Oceans

We benchmarked the inference speed of FlashWeave on the Human Microbiome Project (HMP [33]) and TARA Oceans [34] data sets. For the homogeneous test, we used 2500 oral samples from the HMP data set and measured run time along a gradient of 500, 750 and 1000 randomly selected OTUs (Fig. 2 A). FlashWeave outperformed other methods by factors of 8 to 158 on this task (mean: 67), excluding multiple methods (SpiecEasi-GL, CoNet, mLDM) that did not finish after two days of computation (factor > 339). FlashWeave-S ran on average 33% longer than FlashWeave-F.

On the TARA Oceans data set (289 samples, 3762 OTUs), FlashWeave was on average 29 times faster than the closest follow-up method (SpiecEasi-MB), while all other methods did not finish (factor > 106, Fig. 2 A). FlashWeave-S required 53% more runtime than FlashWeave-F on this benchmark.

Next, we tested the speed of FlashWeaveHE on all five body sites of the HMP data set (5514 samples, 1521 OTUs). FlashWeaveHE-F was 51 times faster than the closest non-FlashWeave follow-up method (SpiecEasi-MB) in this test and on average 371 times faster than standard FlashWeave (other methods did not finish; factor > 518) (Fig. 3 C). FlashWeaveHE-S required 116% more runtime than FlashWeaveHE-F.

To further test its computational scalability, we furthermore used FlashWeaveHE-F to infer a massive ecological network from 504 647 sequencing samples spanning various habitats and conditions, mapped to 75 516 OTUs (98% 16S rRNA identity). The full network finished in 1d10h46min on a High Performance Computing cluster (200 CPU cores).

### Influence of meta variables in the Human Microbiome Project

Meta variables (MVs), such as habitats, conditions (e.g. antibiotics usage) and technical factors (e.g. amplicon vs. whole-genome-shotgun sequencing) can lead to spurious associations between OTUs that share association with the same MV. Furthermore, direct associations between MVs and OTUs can be interesting when investigating which OTUs are for instance directly associated to a particular habitat (independent of microbial interaction partners), prefer certain temperatures or are affected by specific sequencing biases.

We investigated the importance of MVs in the HMP data set, explicitly providing all five body sites and two used primer sets (V13 vs. V35) as MVs to FlashWeave. MVs form central hubs in the resulting interaction network with on average 7.4 times larger neighborhoods than OTUs (Fig. S2 B) and 27.6 times higher betweenness centrality, a measure of node importance in the network. MVs furthermore participate in excluding up to 41.7% indirect OTU-OTU interactions (Fig. S2 A) while constituting only 0.4% of all variables.

However, overall numbers of OTU-OTU interactions increased only moderately (up to 12% across modes) when MVs were omitted, indicating that FlashWeave was generally able to replace MVs in conditional sets with highly associated OTUs in most cases. Accordingly, we found the weak association between shared primer bias and interaction probability (Pearson’s *r* < 0.003) to increase only slightly when omitting primer information (*r* < 0.007), while the univariate network showed a noticeably stronger association (*r* < 0.057). Nonetheless, when only comparing the direct neighborhoods of MVs, we detected 13% - 294% additional associations across modes between OTUs when MVs were not provided (Fig. S2 C), indicating that MV omission may lead to local biases.

### Hidden meta variables and structural zeroes

While MVs may reduce false-positives, information on these variables may not always be available, since rarely all important latent factors are measured and made available in standardized annotation formats. This in particular affects large multi-study data sets which are inherently more heterogeneous in terms of experimental, physicochemical or geographical variables, and usually suffer from inconsistent metadata annotations.

Importantly, artificial correlations caused by MVs manifest themselves partly due to structural zeroes (Fig. 3 A), for example when a data set includes multiple habitats with partially exclusive microbial content or multiple sequencing protocols biased towards disjoint OTU sets. It is therefore desirable for an inference method to show robustness to these structural zeroes in heterogeneous data sets.

In order to compare the robustness of different tools, we computed interaction networks separately for each method and body site in a HMP project data set with increased structural sparsity and then quantified the overlap of the inferred interactions to a network computed on the aggregated data set of all body sites (Fig. 3 B). We find that FlashWeaveHE shows optimal robustness to increased structural zeroes introduced in the cross-site network, with a mean Jaccard overlap between site-specific and cross-site networks of 1.0. In contrast, homogeneous FlashWeave (0.39) and other methods (0.18 - 0.24) are noticeably less robust.

### Inference of a large-scale interaction network from a globally distributed data set of human stool samples

We applied FlashWeave on a data set of 69 818 globally distributed human feces samples (“Global Gut”, GG) from the NCBI Sequence Read Archive database (SRA [23]), spanning 488 studies and multiple sequencing protocols. We extracted protocol information alongside other clustered metadata keywords from SRA annotations and provided these as 96 MVs alongside 10 624 OTUs (98% identity) to FlashWeaveHE-F.

Inference took 3h53min on 20 CPU cores on an Intel Xeon E7-4870 machine (2.4 GHz). The method inferred 30 342 significant interactions between OTUs (after conditioning search) and 13 151 significant interactions between OTUs and MVs (30%). In contrast, the univariate version of the GG network featured strongly increased density at 1 056 262 significant edges, a factor 35 increase. When breaking associations in the GG network via repeated shuffling [34], FlashWeaveHE furthermore reported no false positive interactions.

Reducing the full GG data set to a subset of 8 897 samples corresponding to the American Gut project (AGP [31]) yielded an edge number decrease of 81% (Fig. 4 D), in 57% of which at least one interaction partner was absent in the AGP data set. Notably, these OTUs also showed 90% decreased mean prevalence in GG, thereby representing the rare ecosphere of the human gut. However, OTUs present in AGP also benefited from increased sample size, as 43% additional edges connected these more common microbes.

**Fig. 4.**
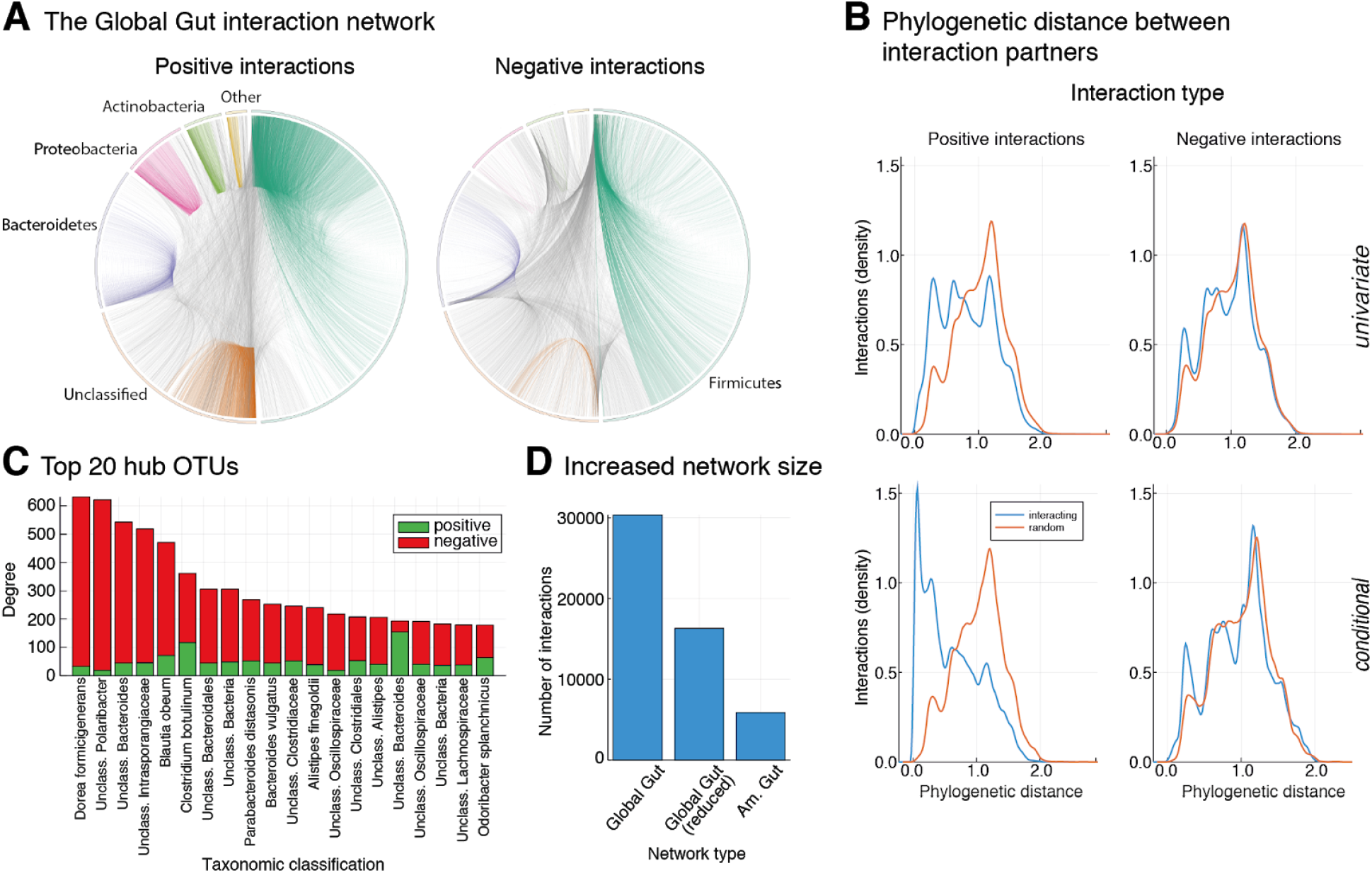
Inference of a large-scale, globally distributed human gut interaction network. High-level overview of positive and negative interactions in the Global Gut network. OTUs are grouped by phylum. Interactions within the same phylum bear that phylum’s color, between-phylum edges are grey. **B** Phylogenetic assortativity pattern for positive interactions and negative interactions. Phylogenetic distance is the summed branch length between interaction partners on a tree of 92 659 OTU representatives (98% 16S rRNA identity). The top panel depicts distributions from the univariate and the lower panel from the conditional Global Gut network. **C** Top 20 OTUs by number of direct interaction partners. “Unclass.” OTUs were not confidently classifiable at species level. **D** Comparison of the number of edges between the full Global Gut network, the Global Gut network reduced to OTUs found in the American Gut Project subset and the American Gut Project network.

The OTU-OTU sub-network is strongly structured (modularity 0.25), indicating the presence of distinct communities. The 20 largest clusters have on average 45 members (up to 89) and feature almost exclusively positive edges between members (mean 99.6%), but only 37.1% - 79.8% (mean 63.3%) positive edges to non-member OTUs.

We found the majority of within-phylum interactions to be positive (47% in *Actinobacteria* up to 75% in *Firmicutes,* mean 64%), while interactions between different phyla were frequently negative (44% in *Firmicutes* up to 95% in *Actinobacteria,* mean 76%). For *Actinobacteria,* which had the highest fraction of negative edges to other phyla, the majority targeted *Firmicutes* (40%) and *Bacteroides* (30%).

Many negative interactions in the GG network were conveyed by few members per phylum (Fig. 4 A, C), constituting negative hubs that were not explainable by our set of MVs (see Methods). These include several species implied in inflammation and disease (*Dorea formicigenerans [35], Clostridium botulinum [36], Odoribacter splanchnicus [37], Bacteroides vulgatus [38]).* Additionally, we found negative associations between multiple *Blautia* OTUs and a *Clostridium difficile* OTU, consistent with previous reports [16, 39].

Phylogenetic assortativity (PA), i.e. increased probability of interaction between evolutionarily more closely related partners, is a repeatedly observed pattern of potential biological interest [14, 15, 40]. We found strong PA in GG for positive edges, while negative edges were closer to the empirical null distribution (Fig. 4 B, lower row). Differences were significant in both cases (p-value < 0.01, two-sample Kolmogorov-Smirnov test), however effect size was 10x increased for positive edges. In contrast, positive edges in the univariate network showed noticeably smaller effect size increase (3.7x, Fig. 4 B, upper row), indicating selective enrichment of this ecologically noteworthy pattern in direct associations.

Among OTUs with the largest numbers of positive neighbors (Fig. S3), potentially constituting keystone species [41], we observed a number of OTUs from *Bacteroides* (genus) and numerous *Clostridiales* (order) OTUs, both known to harbor important cross-feeders in the human gut [42, 43]. Intriguingly, 85% of the top 20 positive hubs were taxonomically uncharacterized at the genus level and 10% even at the phylum level.

Known dependencies between H_2_ producing and consuming microbes have been described in the human gut [44] and indeed we found significantly more positive interactions between H_2_ producers and consumers in GG than in random networks (4.6x increase, empirical p-value < 0.01). This effect was again noticeably weaker for univariate networks (1.8x, p-value < 0.01).

## Discussion

FlashWeave is the first ecological network inference approach that combines i) the estimation of direct interactions, ii) the ability to scale to large-scale data sets with tens of thousands of OTUs and hundreds of thousands of samples and iii) the incorporation of meta variable (MV) information.

We show that FlashWeave typically outperforms state-of-the-art methods by multiple orders of magnitude in terms of run time, compares favourably at recovering gold-standard literature interactions in the TARA Oceans data set and generally surpasses other methods in terms of edge accuracy on a wide selection of synthetic benchmarks. Notably, network recovery generally continued improving with additional samples on most data sets, indicating that FlashWeave benefits from the statistical power of larger data set sizes that are becoming increasingly available.

Sequencing samples do not only increase in number, but also in heterogeneity, as more habitats are being sampled under a plethora of different conditions and with varying experimental protocols. These factors may confound association signals, resulting in biased interaction networks. FlashWeave offers two ways to tackle this challenge: firstly, it features a specialized heterogeneous data mode, FlashWeaveHE, which we find to achieve strongly improved consistency, edge accuracy and run time compared to other methods in the presence of structural zeros which are a typical property of heterogeneous data sets. Secondly, FlashWeave can use MVs to exclude spurious OTU-OTU associations. Exemplified by primers and body sites in the HMP data set, we observed that omission of MVs resulted in noticeable increases of edge density in their neighborhoods, potentially analogous to spurious edges around keystone taxa [41]. Interestingly, we found FlashWeave to be remarkably robust to omission of MVs when looking at the whole network, indicating that overall accurate networks may be inferable even when not all relevant MVs are measured. However, this observation requires further validation.

In addition to helping remove false positive interactions, direct relationships between OTUs and MVs reported by FlashWeave can also yield further insights into the impact of non-microbial factors on study systems. Exemplified by the HMP, we found MVs to be central nodes of the network with many directly associated OTU neighbors, confirming an expected high dependence of many microbes on specific habitats [33] and indicating noticeable primer biases [45].

We demonstrated that FlashWeave scales to modern heterogeneous, large-scale data sets as exemplified by two aggregated cross-study data sets with 69 818 human feces samples (Global Gut) and 504 647 multi-habitat samples that were inferred in less than 4h and 1.5d, respectively. We found that Global Gut strongly improved the number of recovered edges compared to the subset of samples corresponding to American Gut Project data set, indicating that current single-study data sets may miss large fractions of co-occurrence trends due to lack of statistical power and/or heterogeneity. Around half of the missed interactions involved low-prevalence OTUs, highlighting that large-scale data sets such as Global Gut may allow first glimpses at ecological interactions of the hitherto underexplored rare microbiota [21, 46]. This is a crucial advancement, as current analyses are typically restricted to highly prevalent OTUs due to lack of statistical power or computational restrictions.

Encouragingly, the Global Gut network further revealed consistency with several expected biological patterns, such as a known dependency between H_2_ producers and consumers, mutualist hubs mapping to taxa associated with cross-feeding, a previously reported negative interaction partner of *Clostridium difficile,* and phylogenetic assortativity of interacting microbes. The Global Gut network features several hub OTUs with mainly negative interactions, some of which map to disease or dysbiosis-associated species. While we cannot fully rule out potential indirect influences of host-related factors, these hubs nonetheless provide a valuable list of candidates for further experimental validation. In particular, elucidating potential ecological mechanisms (e.g. wide-spread competitive repression or antagonism) driving the negative impact that these OTUs may have on the gut ecosystem would be intriguing.

We further found that the majority of the strongest positive hubs were not classifiable at the genus level, and some even at the phylum level, indicating a crucial knowledge gap among what may be emerging as some of the most important mutualists in the human gut. A remarkable number of these are assigned to the order *Clostridiales,* indicating that positive feedback of this unclassified OTUs from this order on ecosystem maintenance may be more pronounced than currently appreciated [43].

We found clear advantages of using conditioning search in the Global Gut network: both the phylogenetic assortativity and the H_2_ producer/consumer signals were noticeably more pronounced in the conditional network compared to the univariate network. Additionally, the latter included drastically more edges (factor > 35), mirroring results from our synthetic benchmarks and suggesting strong increases in false positives in the absence of conditioning. These observations highlight the advantages of using direct, rather than indirect, associations for interaction prediction, especially for data sets with sufficient power to support this approach. In addition, the unusually strong phylogenetic assortativity signal we observed for positive edges is intriguing and points towards an underappreciated importance of mutualistic or commensal relationships in groups with increased evolutionary coherence. This would suggest that kin selection [47], as previously observed for instance in biofilms [48] or iron acquisition [49], may be more pronounced in the human gut than previously thought. We however note that, despite extensive efforts to remove shared-niche signals, niche-effect contributions to this signal cannot be completely ruled out, necessitating future confirmatory research.

Current limitations of FlashWeave include the handling structural zeros by conservatively discarding any absences. While we found the resulting power reduction to be noticeable for data sets with fewer samples (TARA Oceans), this effect diminished with larger sample sizes in our synthetic benchmarks. Since globally distributed cross-study data sets include even more data, we are thus confident that this approach nonetheless leads to reasonable network predictions. If necessary, more refined models assigning confidences to absences may furthermore be added in the future. As another caveat, we only explored the impact of MVs on the well-researched HMP data set, in future work it would be interesting to investigate the effects of MVs in other contexts, such as marine physicochemical factors or in disease modeling.

The LGL framework is highly flexible, permitting several straightforward extensions, such as more powerful tests [50, 51] and more importantly the prediction of edge directionality in the future. The latter is an exciting prospect that would allow a more causal interpretation of data-driven ecological models and pave the path towards efficient learning of fully predictive models. In the future, such data-driven models could allow to predict the ecological impact of perturbations and catalyze ecological engineering applications.

## Methods

### The Algorithm

FlashWeave is implemented in the Julia Programming Language [52] and based on the Local-to-Global Learning framework (LGL) proposed by Aliferis et al. [24]. Algorithms from this class of causal inference methods start by performing a locally optimal Markov blanket search in order to infer all directly associated neighbors of a target variable *T* (OTU or meta variable in the case of FlashWeave), representing the set of direct causes and effects of *T*. Then, individual neighborhoods are connected through a combinator rule (by default the OR rule in FlashWeave) to form a global association graph. In the final step, not yet implemented in FlashWeave, this undirected skeleton of conditional dependence relationships can then be used as a scaffold to efficiently infer edge directionality and provide further insights into the system of study. In line with results in [53], FlashWeave employs a False Discovery Rate (FDR) adjustment step and omits the costly steps of spouse identification and symmetry correction.

LGL can be instantiated with a wide range of algorithms and conditional independence tests. FlashWeave currently defaults to the successful semi-interleaved HITON-PC algorithm [24] and provides the choice of either discretized mutual information tests (more coarse grained and usually quicker; “fast” mode) or partial correlation tests (more sensitive and usually slower; “sensitive” mode) (Fig. 1 C, Fig. S1).

FlashWeaveHE further specializes these tests to treat zero elements differently (Fig. 1 C, Fig. S1). It makes the general assumption that most zeroes in large, heterogeneous data sets are structural (for instance due to primer or sequencing depth biases, as well as habitat or condition-specific effects) and thereby only considers samples in which both OTUs have a non-zero abundance as reliable for association prediction. Notably, this restriction only concerns the two partners currently tested: OTUs found in the conditioning set retain their absences. This procedure is chosen i) to not discard too much information concerning the tested partners and ii) because uninformative absences, while slightly decreasing power, will otherwise have no or minimal impact on exclusion decisions. While the FlashWeaveHE approach potentially discards some valid absence information and thereby can be less sensitive than vanilla FlashWeave, we see this loss in sensitivity small on heterogeneous data sets with large sample sizes, furthermore counteracted by strongly increased precision in benchmarks (Fig. 3 B, D) and much-decreased runtimes (Fig. 3 C).

Normalization, which accounts for test-specific intricacies and compositionality effects, differs depending on the test type, see Text S1 for details. A discussion of novel heuristics in FlashWeave can also be found in Text S1.

### Accuracy benchmarks

For the NorTA (American Gut) benchmark, synthetic data was generated as described in [15] using the “cluster” and “scale free” topologies, the “amgut.filt” data set and defaults for all other parameters. In order to increase compositionality, we downsampled each sample to depths randomly picked from “amgut.filt”.

For the Ecological Models benchmark, data sets were generated as described in [32], restricted to linear ecological relationships. Three independent data sets (500, 1000 and 2000 samples) were generated per table. To create data sets with multiple disjoint habitats (Fig. 3 D), the three differently sized data sets per table were aggregated with OTUs and interactions assumed as distinct, i.e. with each OTU and interaction only present in one habitat.

For the niche robustness benchmark, we reduced all body sites in the HMP data set to the same number of samples via random subsampling, resulting in 312 samples per site. For each body site, we then picked all OTUs found in at least 10 samples of that site (175 - 619 OTUs) and removed their non-zero counts from all samples from other body sites. The resulting body site-specific data sets were then aggregated into a single table. Inference tools were applied to i) each individual body site table separately, ii) the aggregated data set of all body sites. Finally, the edge overlaps between all sub-networks and the aggregated network were compared using the Jaccard similarity index.

For all benchmarks, parameters reported in Table S1 were used to run each tool.

**Table S1.** Parameters- and versions used for each inference method. All other parameters were kept at their default values.

| Method | Version | Parameters |
| --- | --- | --- |
| FlashWeave | 0.9.0 | max-k: 3, conv: 0.01, feed-forward & fast-elimination enabled |
| SpiecEasi | 0.1.3 | lambda.min.ratio: 1e-2, nlambdas: 15, nrep: 20 |
| eLSA | 81a2ee0 | p: theo, n: percentileZ, f: zero, q-value < 0.05 |
| SparCC | See<br>SpiecEasi | q-value < 0.05, FDR correction via p.adjust (option "BH") in R [54] |
| CoNet | 1.1.1 | methods:<br>correl_spearman/dist_kullbackleibler/dist_bray,<br>multitestcorr: benjaminihochberg,<br>q-value < 0.05 |
| mLDM | 1.0 | Z_mean: 1 |

### Literature interaction predictions

To reduce computation time, we filtered the TARA Oceans data set for OTUs participating in at least one genus-level literature interaction reported in [34]. After removing samples with no reads, the final data set consisted of 234 samples and 702 OTUs. Edges predicted by each tool were sorted according to reported weights (merged q-value in the case of CoNet, which uses multiple weight measures, Pearson’s *r* for SparCC) and this ranking was plotted as cumulative curves (Fig. 2B).

### Speed benchmarks

The HMP data set consisted of 5514 samples from the body sites Oral, Gastrointestinal tract, Urogenital tract, Skin and Airways. Samples were mapped to OTUs at 96% 16S rRNA identity using MAPseq (version v1.0 [55], confidence > 0.5) and the full-length 16S reference provided with MAPseq. The data set was further filtered for OTUs present in > 20 samples.

For the TARA Oceans data set, we used the OTU counts table provided by [34]. After filtering for OTUs present in > 50 samples and samples with at least one read, the data set contained 289 samples and 3762 OTUs.

Parameters for each network inference tool were as reported in Table S1. Since not all tools readily support parallelism, all benchmarks were conducted on a single core on an AMD Opteron 2347 HE machine (1 GHz).

### Meta variable analysis in the HMP

An MV was counted as explaining an association if it was present in the set of conditional variables that lead to that association’s exclusion (Fig. S2 A). The association between shared primer influence and interaction probability was estimated by computing, for each pair of OTUs (*O_i_, O_j_*), the absolute difference of association strengths in the HMP network between *O_i_* and the primer MV and *O_j_* and the primer MV, leading to small values for OTU pairs with similar primer influence and larger values for differences in influence. These values were then correlated (Pearson’s *r*) with the interactions strength’s between each *O_i_* and *O*_j_.

### Global Gut network analysis

#### Data set creation

Studies from the NCBI Sequence Read Archive database (SRA [23]) were filtered for human samples through automated parsing of metadata annotation fields, matching at least one of the following rules: 1) “Human” or “Homo sapiens” is found in the host name field, 2) “9606” is found in either the host taxon ID or sample taxon ID field, or 3) “human <*> metagenome” is found in the organism field, where “<*>“ is a wildcard for either “gut” or “gastrointestinal”. For matching samples, a list of keywords was parsed from all main annotation fields and further curated to remove uninformative terms, resulting in a set of keywords assigned to each sample. Samples were then further filtered for gut association through the keywords “intestinal”, “intestine”, “alimentary”, “bowel”, “cecum”, “crohn”, “gut”, “colon”, “commensal-gut”, “diarrhoea”, “digestive-tract”, “digestive tract”, “duodenum”, “enteric”, “enteritis”, “enterocolitis”, “enteropathogenic”, “enterohemorrhagic”, “equol”, “feces”, “gastroenteritis”, “gastrointestinal”, “ileum”, “ileostomy”, “jejunum”, “meconium”, “mesentery”, “mid-gut”, “probiotic”, “rectum”, “stec”, “vibriosis”. Keywords of these samples were further checked for terms not related to gut, followed by manual review of matching samples via the SRA web service and removal in case of non-gut origin.

The final set of samples was downloaded and mapped to OTUs at 98% 16S rRNA identity using MAPseq (version v1.0 [55], confidence > 0.5) and the full-length 16S reference provided with MAPseq (hierarchically clustered with HPC-CLUST [56]; average linkage). We removed samples with less than 100 mapped reads and OTUs found in less than 200 samples (see Table S2 for SRA accessions of the final sample set). Taxonomy was assigned to OTUs based on a 90% consensus over the full taxonomic lineages of all OTU member sequences. For sequences belonging to RefSeq [57] genomes or culture collection strains, the annotated taxonomy as provided by NCBI (accessed in December 2017) was used. The remaining sequences were taxonomically classified through mapping onto the RefSeq set with MAPseq (confidence > 0.5).

In addition, we retrieved sequencing method information from the SRA (“WGS”, “AMPLICON” “RNA-SEQ” or “OTHER”) and filtered the previously extracted metadata keywords for a set of 128 potentially interesting terms such as “fibre”, “antibiotics” and “cancer”. This metadata information was used to create a MV table which was further hierarchically clustered into 96 MV groups (average linkage, unweighted Jaccard similarity > 0.9). See Table S3 for representatives used for each group.

The OTU table and the MV group table were finally used as input to FlashWeaveHE-F with parameters from Table S1.

#### FDR estimation and modularity

For the false positive analysis, we generated a null model by breaking associations between taxa through sequencing depth-conserving shuffling of the GG dataset (see [34]). Modularity [58] was computed based on cluster assignments from Markov Chain Clustering (MCL [59] version 14-137, inflation parameter 1.5).

#### Impact of meta variables on negative hubs

In order to estimate whether negative associations of the top 20 negative hub OTUs could be explained by MVs, we collected all negative associations of these OTUs and computed for each MV, how often samples assigned with this MV contributed to a positive or negative association signal within the negative edges. We then compared MV frequencies of negative contributions to those of positive contributions and found no significant difference (p-value > 0.99, paired T-test), indicating that positive and negative association signals were overall driven by samples with highly similar MV distributions.

#### Phylogenetic assortativity

For phylogenetic tree construction, the alignment of representatives of all 98% 16S rRNA identity OTUs in the MAPseq reference database (92 659 full-length 16S rRNA sequences), created with INFERNAL (version 1.1.2 [60]) and microbial secondary structure model SSU-ALIGN [61], was used. The phylogenetic tree was then reconstructed using fasttree (version 2.1.3 [62]) with the GTR substitution model and otherwise default options. For the phylogenetic assortativity analysis, the Global Gut network was reduced into two separate networks restricted to edges and vertices participating in only positive and negative associations, respectively. To generate a random background, vertices in each graph were randomly connected to create a network with vertex and edge numbers matching the original graph. Phylogenetic distance between interaction partners was calculated as total branch length between the corresponding leaves.

#### Associations between H_2_ producers and consumers

OTUs mapping to H_2_ producing and consuming taxa (taken from [44]) were identified in the Global Gut network. The number of positive interactions between these groups was then compared to those found in 100 randomly generated networks with conserved expected positive degree for each OTU.

#### Normalization comparison

The subset of Gastrointestinal tract samples from the HMP data set was filtered along sequencing depth and OTU prevalence gradients, followed by normalization with *clr* (pseudo-count 1) and *clr-adapt* normalization (Text S1). Associations were inferred using FlashWeave-S with max-k 0 (univariate) and max-k 3 (conditional) and all other options as in Table S1. For the oral comparison, the oral subset of the HMP data set (1000 OTUs) was used and network inference was done with 20 CPU cores.

### Data and Software Availability

FlashWeave is implemented in Julia [52] and freely available as open source from https://github.com/meringlab/FlashWeave.jl.

## Supplementary methods and results

### Novel heuristics

Learning the direct neighborhood of target variable *T* with the si-HITON-PC algorithm has a run time complexity of *O*(|*V*|2^|*PC*(*T*)|^) [24], where *V* are variables in the system and *PC*(*T*) is the Parent-Children set of *T*, i.e. the set of its directly associated neighbors (or the Markov blanket *MB*(*T*) minus spouses [24]). Runtime thus depends linearly on the number of variables and exponentially on the size of the direct neighborhood.

FlashWeave implements all options and algorithmic shortcuts suggested by Aliferis et al. [24] (max-k heuristic, h-ps reliability criterion, FDR correction, optimal variable ordering). In order to achieve the speed reported in this study, we furthermore extended the original algorithm through a number of additional high-performance heuristics that we will explain in more detail below.

The first novel algorithmic shortcut we term *feed-forward* heuristic, which is a parallel variation of traditional backtracking [63]. The key observation for this heuristic is that the size of individual neighborhoods can vary substantially in scale-free networks, such as microbial co-occurrence networks ([10] and citations therein), with exponential impact on runtime. From the scale-free property follows that keystone species *A* (with many dependent neighbors) will have a large number of neighbors *B* that themselves are not keystone species (few neighbors) and are thus considerably quicker to compute. Now, for the edge *A → B* to be included in the final, global network by means of the OR combinator rule, it is sufficient if the reverse direct link *B → A* is proven, which takes exponentially less time to compute. *feed-forward* exploits this by prioritizing computation of variables expected to have smaller neighborhoods (as approximated by their univariate neighborhood size) and relaying the information of detected direct links to computationally more intensive variables. If during the computation of the neighborhood of *A* the next variable *B* to be tested was already shown to be a neighbor, it automatically enters the set of neighbor candidates of *A* without formally performing all tests. In contrast to vanilla backtracking, *feed-forward* is applied in a parallel computation setting, where candidate lists of all nodes are periodically updated with the latest information from other neighborhoods as it becomes available, allowing the most expensive nodes to leverage a maximum amount of information to cut down runtime.

The second computational shortcut we term *fast-elimination* heuristic. A large amount of time can be spend in the final elimination phase of si-HITON-PC, in which all previously skipped tests between candidates passing the interleaving phase are performed. In the original algorithm, even if a variable *S* is discarded during the elimination phase, it will still be included in future conditioning sets, thereby inflating the number of conducted conditional independence tests. If many variables are formally discarded during the elimination phase, but still included in future conditioning sets, the result can be an exponential increase in necessary tests, making the elimination phase particularly costly. *fast-elimination* addresses this computational hurdle by not considering a removed variable *S* for any subsequent conditioning sets. An intuitive motivation for this approach is that, if a variable *S* was shown earlier to not be part of the neighborhood of *T*, it should also not be required to render further candidates independent of *T* as it’s not part of its Parent-Children set.

As another shortcut, we implemented a convergence criterion that periodically checks whether links in the network still show substantial changes over time. If the network has reached convergence, all remaining candidates are assumed to be conditionally independent of their target variables. This criterion is based on our observation that the naive algorithm can stall on single nodes with large neighborhoods due to the exponential runtime dependency of si-HITON-PC on neighborhood size. However, candidates still to be checked at this point tend to be weak, since they i) appear late in the relevance-sorted candidate list and ii) have been proven to not be neighbors in the reverse direction (otherwise the *feed-forward* heuristic would apply). We observe that the majority of these late links are thus finally discarded after substantial computational effort, with minimal effect on network structure. While using this type of convergence threshold may in theory lead to biased edge omissions since it selectively bypasses computation in variables with large neighborhoods, we didn’t detect meaningful biases of this kind in the networks we tested.

As a final option, FlashWeave can be instructed to run only up to a certain (by default large) number of tests per node, assuming that performing such a high number of tests provides reasonable safety that the current candidate will not be discarded by additional tests. This effectively puts an upper bound on the exponential behaviour of si-HITON-PC and helps to prevent extensive run times on single variables with large neighborhoods, with empirically minimal effect on network structure. However, FlashWeave will flag these interactions and warn the user in case the boundary is breached.

### Normalization

Sequencing data is subject to mainly technically determined and thus arbitrary variations in sequencing depth, making it compositional in nature. Compositionality impedes naive correlation analysis without adequate correction [29, 30]. Common approaches to properly analyze compositional data include various log-ratio based approaches, such as log-ratio transformations [29].

Similar to SpiecEasi [15], FlashWeave uses the centered log-ratio (*clr [29]*) approach for compositionality correction of a vector *x* of compositional microbial abundances:

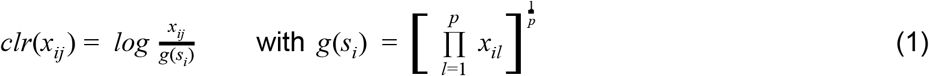

where *g*(*s_i_*) describes the geometric mean of all compositional abundances in sample *s_i_*, *p* the total number of OTUs and *clr*(*x_ij_*) the *clr*-transformed value of the compositional abundance of microbe *j* in sample *s_i_*.

An inherent shortcoming of logarithm-based methods is the handling of absences (zeroes) in the input data. This is usually circumvented by applying a fixed pseudocount (for example 1) to the input data which then allows proper computation of logarithms. Our analyses reveal that this approach can work decently well for strongly filtered and depth-homogeneous data sets, but introduces noticeable biases when applied to data sets that include rare OTUs and/or samples with particularly low sequencing depths (Fig. S4 A, left column; Fig. S4 B). In such data, we observe extensive increases in univariate network density, which renders the subsequent conditioning search in FlashWeave unusually slow (Fig. S4 C). Importantly, most of these additional univariate associations are removed during conditioning search (Fig. S4 A, left column), indicating their spurious nature.

To be more precise, absences of comparatively rare OTUs in low-depth samples can, after *clr* transformation, become values higher than the OTU’s mean *clr* -transformed abundance across all samples, while absences in high-depth samples result in transformed values below these OTU’s means. This depth-based deviation from the mean results in the observed artificial correlation signal and notably is driven solely by applying the same fixed pseudo-count both to low-depth and high-depth samples. While homogenizing sequencing depth through sample removal and filtering of rare OTUs reduces this signal (Fig. S4 A, left column), large amounts of valuable data are potentially removed by this approach.

As an alternative method to reduce the pseudo-count driven association signal, we suggest a modification to classic fixed pseudo-counts, which we term “adaptive pseudo-counts”, resulting in the normalization scheme *clr-adapt.* In this approach, initially a fixed pseudo-count *π_max_* is applied to the sample with the highest sequencing depth (*s_max_*). Then solving

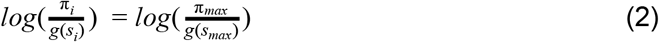

for π_i_ (the adaptive pseudocount for sample *s_i_*) leads to

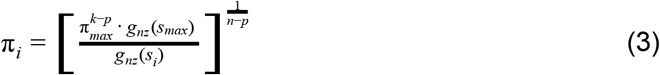

where *g_nz_*(*s*) is the geometric mean of all non-zero abundances in sample *s*, *k* is the number of absences in sample *s_max_* and *p* is the number of OTUs. Formula 3 is applied to all samples besides *s_max_* in order to determine sample-specific adaptive pseudo-counts. These are then applied to their respective samples, followed by usual *clr* transformation (formula 1). This results in the same transformed absence counts in all samples and ensures that all absences are below each OTU’s mean *clr* -abundance, which avoids bi-directional pseudo-count driven deviations from the mean. Using this approach, we observe strongly reduced univariate network densities, discard fractions and run times (Fig. S4).

FlashWeaveHE (f) also utilizes *clr* transformation for compositionality correction, albeit slightly modified. Since FlashWeaveHE does not include absences into its association calculations (see Methods), it does not transform zeroes and and thus requires no (adaptive) pseudo-counts. Instead, only non-zero compositional abundances are used to compute the compositional center (geometric mean, formula 1) and then transformed, resulting in the normalization scheme *clr-nonzero.*

The (f) modes of FlashWeave and FlashWeaveHE differ from the (s) modes by applying mutual information tests which necessitate data discretization. FlashWeave (f) uses a straight-forward discretization scheme: all non-zero abundance values become one, while absences remain zero. This approach makes *clr* normalization and pseudo-counts unnecessary and is inherently robust to compositional artifacts. It is furthermore less affected by sequencing depth differences, since depth only affects OTU-presence and absence but not abundance. FlashWeaveHE (f), on the other hand, discretizes all *clr-nonzero* transformed values into two bins per OTU (“high” abundance vs “low” abundance), with bins separated by the median.

Meta variables (MVs) are by default not normalized for sensitive FlashWeave(HE) modes and should thus be provided in a sensible pre-normalized form by the user if necessary. For (f) modes, continuous MVs are by default discretized into two bins separated by their median quantity.

**Fig. S1.**
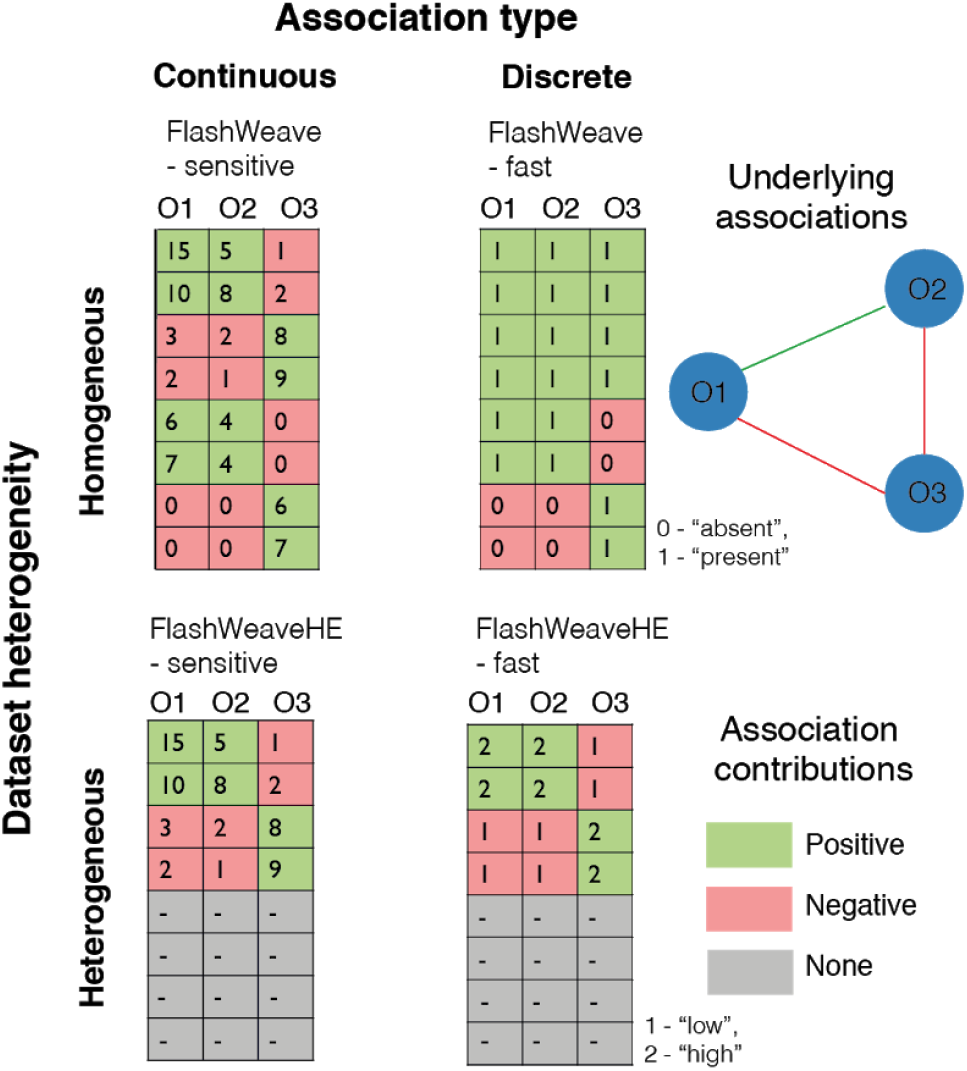
Differences in association inference amongst FlashWeave modes. Sensitive modes (-S) of FlashWeave use full abundance information (“continuous”), while fast modes (-F) work in a discretized fashion. In contrast to FlashWeave, FlashWeaveHE excludes samples where one partner is absent (colored grey). However, it still includes absences of OTUs found in the conditioning set.

**Fig. S2.**
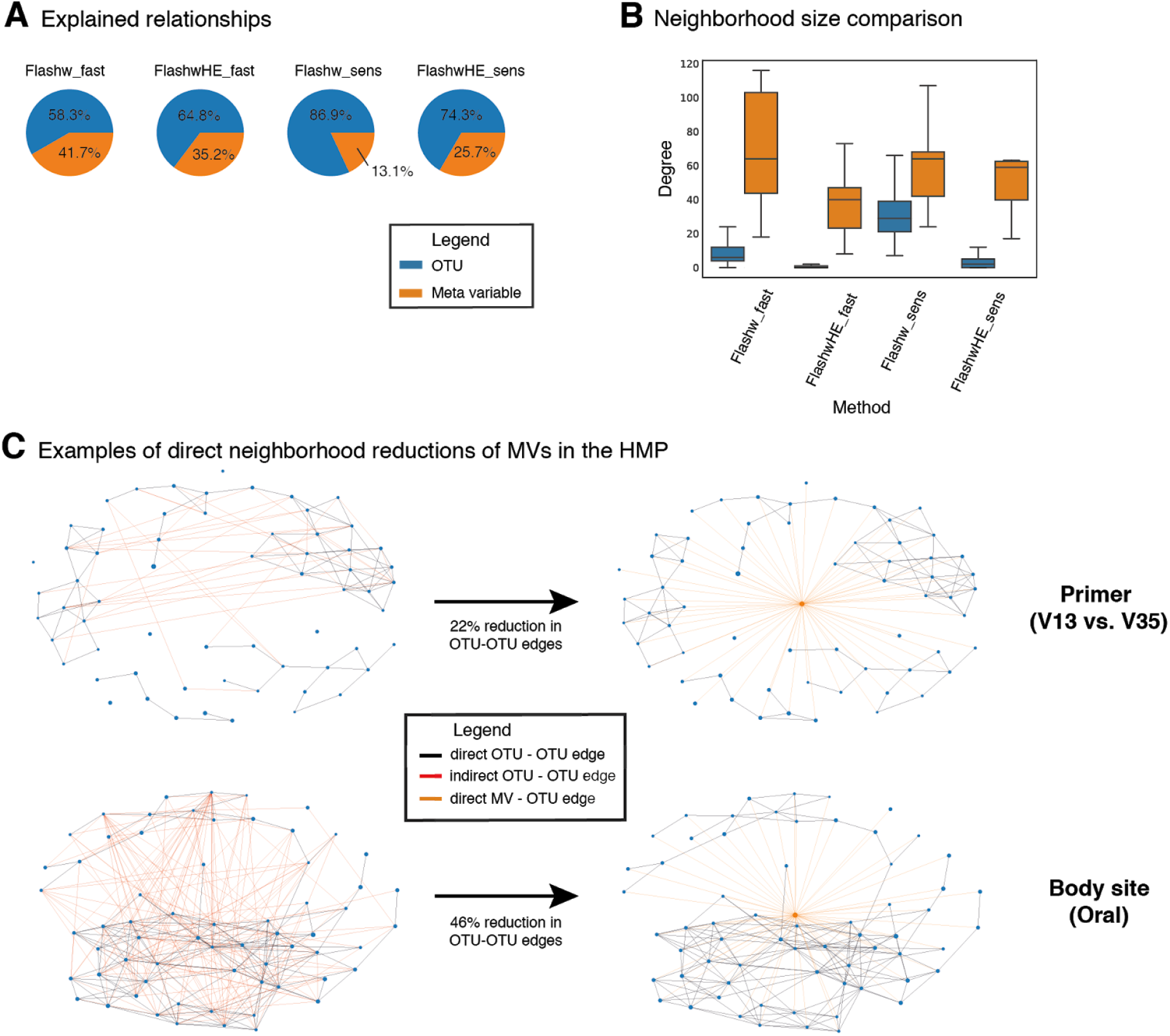
Importance of meta variables (MVs) in the HMP network. **A** Fractions of indirect OTU-OTU associations excluded through at least one MV (orange) or solely through other OTUs (blue). **B** Number of direct neighbors of OTUs and MVs. **C** Exclusion of indirect edges (red) in the direct neighborhoods of the MVs “Primer” and “Oral” (orange edges) in the HMP data set when explicit MV information is provided to FlashWeave-S. Edges retained after including MV information are colored black.

**Fig. S3.**
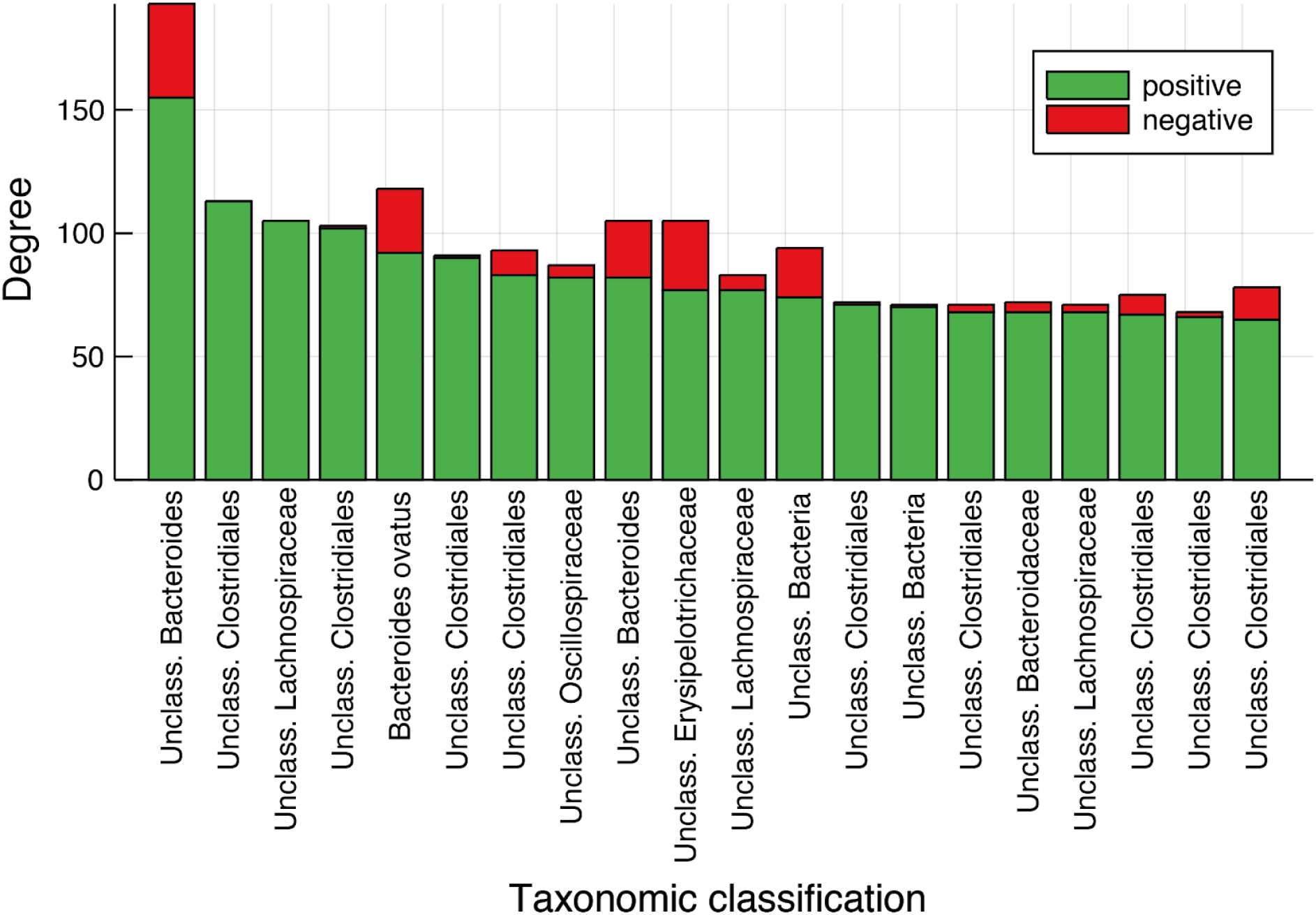
Top 20 OTUs by number of direct positive interaction partners. “Unclass.” OTUs were not confidently classifiable at species level.

**Fig. S4.**
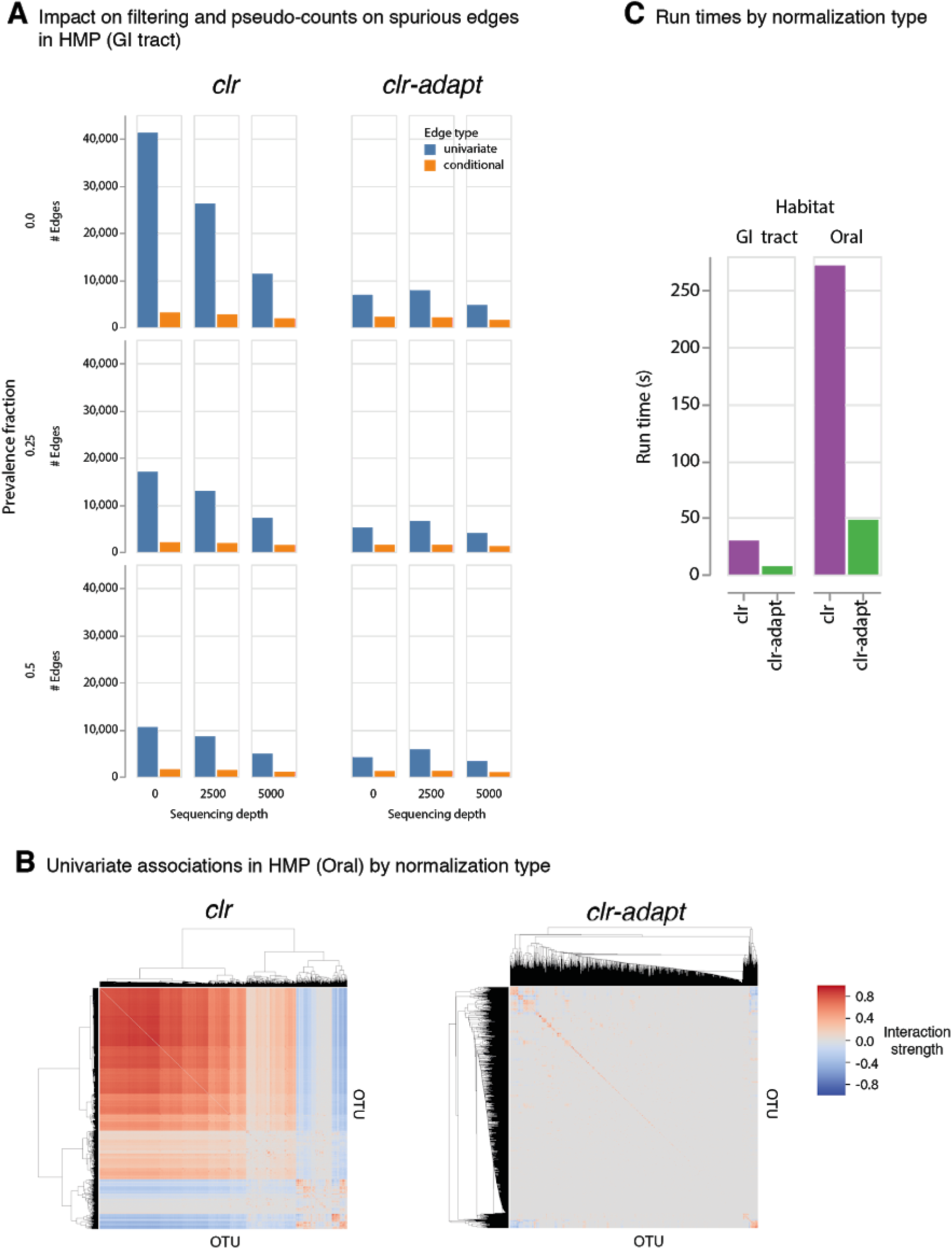
Impact of normalization on network inference. **A** Network sizes along gradients of increasingly strict OTU prevalence (rows) and sequencing depth (columns) filtering of the Gastrointestinal tract subset of the HMP data set, stratified by normalization type (*clr* with pseudo-count 1 vs. *clr-adapt).* **B** Univariate association patterns in HMP Oral (1000 OTUs) by normalization type (no additional filtering). **C** Run time comparison between normalization types on HMP (GI tract) and HMP (Oral).

